# A scalable strategy for high-throughput GFP tagging of endogenous human proteins

**DOI:** 10.1101/055285

**Authors:** Manuel D. Leonetti, Sayaka Sekine, Daichi Kamiyama, Jonathan S. Weissman, Bo Huang

**Author notes:** M.D.L. and S.S. contributed equally to this work. To whom correspondence should be addressed (D.K.) (J.S.W.) (B.H.).

## Abstract

A central challenge of the post-genomic era is to comprehensively characterize the cellular role of the ∼20,000 proteins encoded in the human genome. To systematically study protein function in a native cellular background, libraries of human cell lines expressing proteins tagged with a functional sequence at their endogenous loci would be very valuable. Here, using electroporation of Cas9/sgRNA ribonucleoproteins and taking advantage of a split-GFP system, we describe a scalable method for the robust, scarless and specific tagging of endogenous human genes with GFP. Our approach requires no molecular cloning and allows a large number of cell lines to be processed in parallel. We demonstrate the scalability of our method by targeting 48 human genes and show that the resulting GFP fluorescence correlates with protein expression levels. We next present how our protocols can be easily adapted for the tagging of a given target with GFP repeats, critically enabling the study of low-abundance proteins. Finally, we show that our GFP tagging approach allows the biochemical isolation of native protein complexes for proteomic studies. Together, our results pave the way for the large-scale generation of endogenously tagged human cell lines for the proteome-wide analysis of protein localization and interaction networks in a native cellular context.

**SIGNIFICANCE STATEMENT:** The function of a large fraction of the human proteome still remains poorly characterized. Tagging proteins with a functional sequence is a powerful way to access function, and inserting tags at endogenous genomic loci allows the preservation of a near-native cellular background. To characterize the cellular role of human proteins in a systematic manner and in a native context, we developed a method for tagging endogenous human proteins with GFP that is both rapid and readily applicable at a genome-wide scale. Our approach allows studying both localization and interaction partners of the protein target. Our results pave the way for the large-scale generation of endogenously tagged human cell lines for a systematic functional interrogation of the human proteome.

## INTRODUCTION

More than a decade after the completion of the Human Genome Project (1), over 30% of human genes still lack clear functional annotation (2, 3). Functional tagging is a powerful strategy to characterize the cellular role of proteins. In particular, tags allow access to two key features of protein function: localization (using fluorescent tags) and interaction partners (using epitope tags and immuno-precipitation). Hence, by tagging proteins in a systematic manner, a comprehensive functional description of an organism's proteome can be achieved. The power of systematic tagging approaches is best illustrated by studies conducted in the budding yeast Saccharomyces cerevisiae (4). In particular, a genome-wide collection of GFP-tagged yeast strains enabled the systematic study of protein localization in live cells (5), while libraries of strains expressing TAP epitope-fusion proteins paved the way for the large-scale isolation and proteomic analysis of protein complexes (6, 7). One of the great advantages of yeast genetics (especially in S. cerevisiae) is the efficiency and relative simplicity of PCR-based homologous recombination (8). As a result, functional tags can be easily inserted in a gene locus of interest, preserving endogenous expression levels and minimizing genomic disruption. Together, these genome-wide tagged libraries helped provide a comprehensive snapshot of the yeast protein landscape under near-native conditions (4, 5, 9–11).

The development of CRISPR/Cas9-based methods has profoundly transformed our ability to directly tag human genes at their endogenous loci by facilitating homologous-directed repair (HDR) (12, 13). These methods pave the way for the construction of genome-wide, endogenously tagged libraries of human cells. Any large-scale effort should ideally meet four criteria: (i) scalability, to allow large numbers of genes to be tagged in a time-and cost-effective manner; (ii) specificity, limiting tag insertion to the genomic target (ideally in a “scarless” manner that avoids insertion of irrelevant DNA such as selection marker genes); (iii) versatility of the tag, preferably allowing both localization and proteomic analyses and (iv) selectability of knock-in cells. Recently, a strategy based on electroporation of Cas9/sgRNA ribonucleoprotein complexes (RNPs) has been reported that enables both scalability and specificity (14, 15). In this approach, RNPs are assembled in vitro from purified single-guide RNA (sgRNA) and Cas9, both of which can be obtained commercially or rapidly generated in house. The HDR template containing tag sequence and homology arms to the target locus is supplied as a long single-stranded DNA (ssDNA), commercially available up to 200 nucleotides (nt). Electroporation of RNP and ssDNA donor into cells results in very high (>30%) knock-in efficiencies, while the limited RNP half-life in vivo minimizes off-target integration (14). We reasoned that this strategy would be well suited for large-scale knock-in efforts in human cells, and envisioned that GFP would be a functional tag of choice: on top of being a fluorescent marker, GFP is also a highly efficient purification handle for protein capture and subsequent proteomic analysis (16–18). GFP-tagged cells are also readily selectable by flow cytometry.

Here we present an experimental approach for the functional tagging of endogenous human loci that meets all four of the above criteria. We recently described how a split GFP system allows functional GFP endogenous knock-in using a minimal tagging sequence (GFP11, corresponding to the 11th beta-strand of the super-folder GFP beta-barrel structure) (19). When expressed in the same cell, GFP11 and its complementary GFP fragment (GFP1–10) enable functional GFP tagging upon complementation (20). A key advantage of the GFP11 sequence is its small size (16 amino acids): this allows commercial ssDNA oligomers to be used as HDR donors, circumventing any requirement for molecular cloning. Here we show that electroporation of Cas9 RNPs and GFP11 ssDNA donors in cells constitutively expressing GFP1–10 enables the fast (<1 day) and robust generation of GFP-tagged human cell lines. Tagged proteins are expressed from their endogenous genomic loci with minimal genomic disruption. Applying this strategy to a set of 48 human proteins, we demonstrate the scalability of our method and define the expression threshold for detection of knock-in cells by flow cytometry. We next present how our protocols can be easily adapted to allow the knock-in of GFP11 repeats at a given locus, which critically allows the functional characterization of low-abundance proteins in a native context. Finally, we describe how GFP11 tagging also enables the isolation of endogenous protein complexes for proteomic analysis, highlighting the versatility of our approach to examine complementary aspects of protein function.

## RESULTS

**GFP11 and RNP electroporation enable cloning-free, high efficiency GFP tagging in human cells.** Our approach combines two existing methodologies. First, we took advantage of a split-GFP system that separates the super-folder GFP protein into two fragments: GFP1–10 and GFP11 (20). GFP1–10 (i.e. GFP without the 11th beta-strand) contains an immature GFP chromophore and is non-fluorescent by itself. Upon co-expression in the same cell, GFP1–10 and GFP11 assemble non-covalently and spontaneously reconstitute a functional GFP molecule (20, 21). Fused to a protein of interest, GFP11 recruits its GFP1–10 partner and enables fluorescent tagging by GFP complementation (Fig. 1A). The fluorescent intensity of the complemented GFP11/GFP1–10 complex is essentially identical to that of full-length GFP (19, 21). Second, we used electroporation of preassembled Cas9 RNPs to achieve high-efficiency genome editing in human cells (14, 15). In particular, very high rates of knock-in have been reported using timed delivery of Cas9 RNPs and ssDNA HDR templates in human cell lines (14). A critical advantage of this strategy is that all the components required for editing (Cas9, sgRNA and HDR template) are commercially available or rapidly synthesized in house. Cas9 protein can be readily purified from E.*coli* overexpression cultures (22). Similarly, sgRNAs can be easily transcribed in vitro (14, 23). Purified Cas9 and synthetic sgRNAs can also be obtained commercially. Finally, synthetic ssDNA oligomers are readily available, with a typical size limit of 200 nt. Here, the small size of GFP11 (16 amino acids) is key: 200 nt is enough to include the GFP11 sequence (57 nt, including a 3-amino acid linker) flanked by two ∼70 nt homology arms for HDR. Together, the GFP11 methodology and Cas9 RNP electroporation enable the high-efficiency fluorescent tagging of human proteins at their endogenous loci with minimal preparation. Importantly, no molecular cloning is required.

**Fig. 1.**
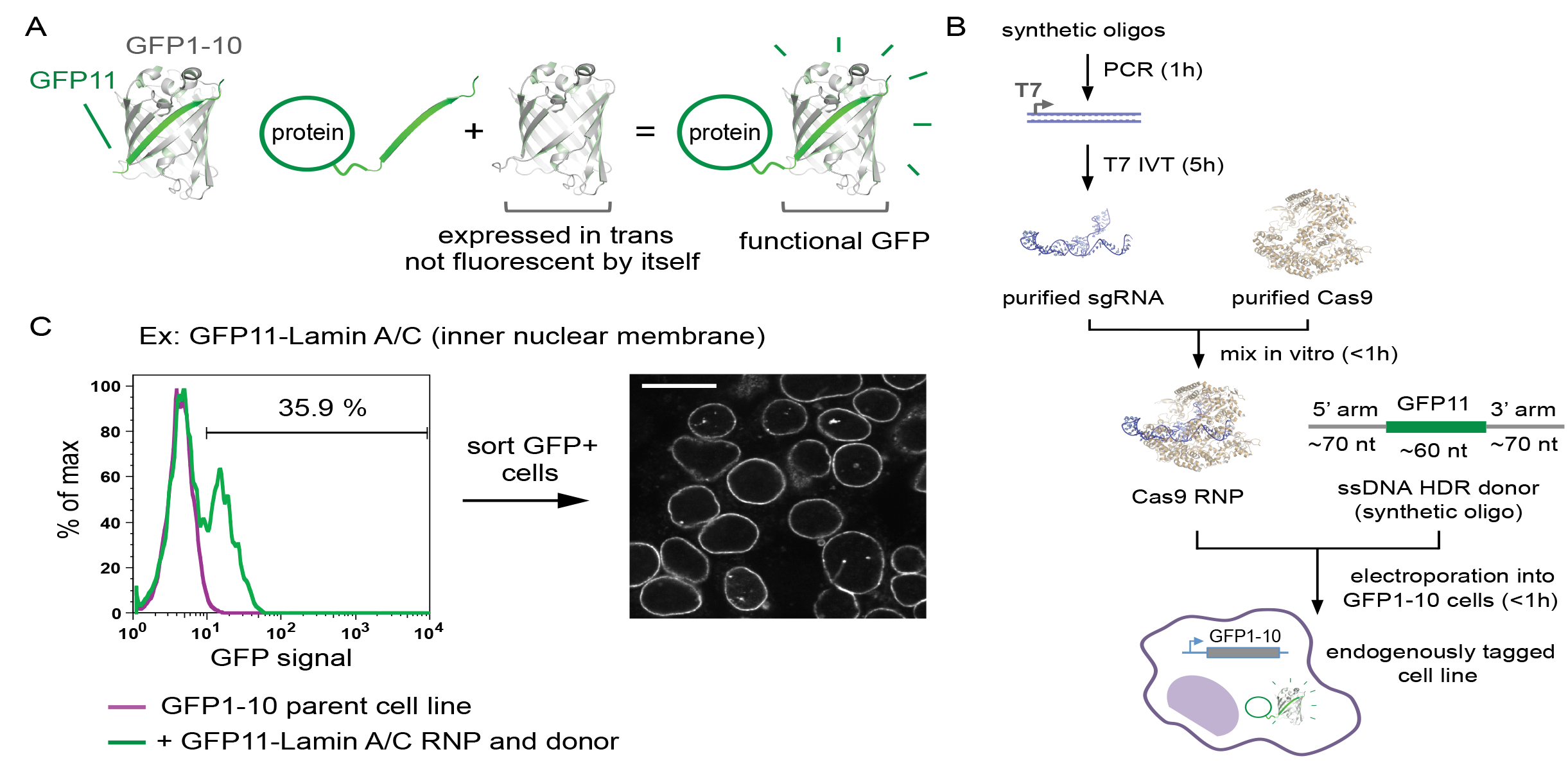
Endogenous GFP11 tagging using Sas9 RNP. (A) Principle of GFP11-mediat-ed tagging. (B) Experimental workflow. T7 IVT: in vitro transcription using T7 polymerase. (C) GFP11 knock-in at the Lamin A/C N-terminus. Knock-in efficiency was analyzed by flow cytometry (left panel, showing the distribution ofGFP fluorescence as histogram plot). GFP-positive cells were isolated by FACS and characterized by confocal microscopy (right panel,showing GFP fluorescence, scale bar =10 μm).

Our experimental design is outlined in Figure 1B. sgRNAs are transcribed in vitro following PCR assembly of a template including a T7 promoter. RNPs are obtained by mixing of sgRNAs with purified Cas9 protein and supplemented with HDR ssDNA donor. Finally, the RNP/donor mix (100 pmol each) is electroporated into cells that constitutively express the GFP1–10 fragment. For all experiments, we used a human 293T^GFP1–10^cell line in which the fragment is stably expressed under the control of a strong SFFV promoter by lentiviral GFP1–10 integration (hereafter, 293T^GFP1–10^). To test our strategy, we targeted the inner nuclear membrane protein GFP1–10 lamin A/C in 293T-cells using an N-terminal GFP11 tag. Flow-cytometry analysis demonstrated very high efficiency of functional GFP tagging (>35%,Fig. 1C). To verify that the GFP signal corresponds to GFP-tagged lamin A/C, we sorted the GFP-positive cells (as a polyclonal population) and analyzed them by microscopy. All cells exhibited a clear GFP localization limited to the immediate peri-nuclear region (Fig. 1C). Low-magnitude images are shown in Figure S1, demonstrating a specific peri-nuclear localization of GFP-tagged lamin A/C across the entire cell population. These results demonstrate that functional tagging with GFP11 is effectively exclusively on-target, eliminating the need to obtain clonal cell lines.

Our protocol can be performed in less than a day (Fig. 1B). We use in-house in vitro transcription as a cost-effective alternative to synthetic sgRNAs, while using commercial synthetic sgRNAs could further shorten the time needed to conduct the experiments. We routinely use column-based methods for sgRNA purification, but solid-phase reversible immobilization (SPRI) magnetic beads can be used to the same effect and are best suited for large-scale preparation in multi-well format (24). The final electroporation step is done in 96-well format so that a large number of cell lines can be processed in parallel. Therefore, our method is well suited for the rapid and robust generation of libraries of GFP-tagged human cell lines in multi-well format. Detailed protocols are available in the Materials and Methods section.

**Library-scale generation of knock-in cell lines.** To test whether our experimental design was applicable to the library-scale generation of endogenously tagged human cell lines, we applied it to a set of 48 human genes in 293T^GFP1–10^ cells. This experiment addresses two complementary questions. First, we wanted to evaluate whether most loci would be amenable to GFP11 knock-in. Second, we sought to determine the threshold of endogenous protein expression that yields a sufficient level of GFP fluorescence for the detection of knock-in cells by flow cytometry or microscopy.

We chose to tag proteins with distinctive sub-cellular localizations so that microscopy analysis of GFP-positive cells would be a good predictor of on-target knock-in. GFP11 was introduced at either N-or C-termini. For each protein target we tested a single sgRNA, selected to induce genomic cleavage within 30 nt of the chosen terminus. HDR donor templates were designed to disrupt the sgRNA recognition site in order to prevent further cleavage of knocked-in sequences by Cas9. Finally, we characterized the efficiency of GFP11 knock-in by flow cytometry (Fig. 2A). Out of the 48 genes we targeted, 30 (i.e. 63%) gave rise to a clear population of GFP-positive cells. For each of these 30 successful targets, we analyzed the resulting cells by confocal microscopy and confirmed that GFP fluorescence matched exclusively the expected subcellular localization of the corresponding protein (Fig. 2A; complete data for all 30 cell lines is shown in Supplementary Figure S2). We further characterized four of these cell lines by fluorescence-activated cell sorting (FACS) followed by immuno-fluorescence using antibodies specific to the target proteins. In all cases, GFP and immuno-fluorescence signals coincided entirely, validating the specificity of GFP11 knock-in (Fig. S3). Altogether, this initial library-scale analysis proves that our method is scalable for the specific endogenous GFP tagging of a large number of human genes.

**Fig. 2.**
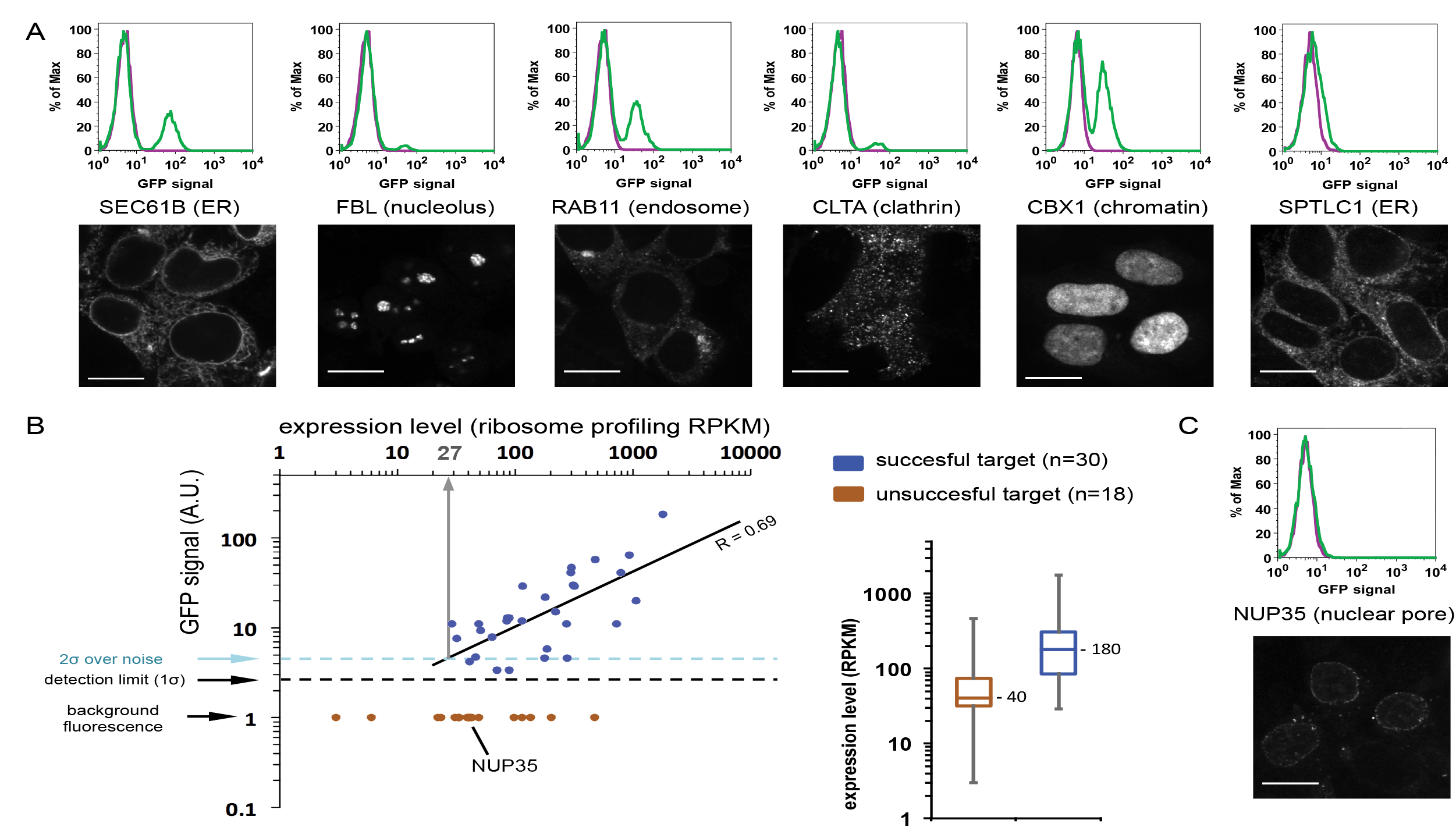
Library-scale GFP11 tagging of 48 different gene targets. (A) Examples of successful targets showing knock-in efficiency (flow cytometry histograms, top panels) and confocal microscopy analysis (GFP fluorescence, bottom panels; scale bar =10 μm). As GFP intensity varies widely across different targets, the different images showed here use different levels of brightness and contrast. (B) Correlation between target expression level (defined as ribosome profiling RPKM) and GFP signal (as measured by flow cytometry, arbitrary units scaled to background fluorescence =1). The 30 successful targets and 18 unsuccessful targets are shown as blue and brown dots, respectively. For successful targets, a linear regression is shown (solid line, Pearson R =0.69). Insert: box plots showing RPKM distribution for unsuccessful vs. successful targets. Boxes represent 25th, 50th and 75th percentiles. Whiskers represent minimum and maximum values. (C) Analysis of NUP35 GFP11 knock-in by flow cytometry (top panel) and confocal microscopy (bottom panel, scale bar =10 μm). NUP35 knock-in cells are not detected by flow cytometry but can be identified by microscopy.

To test the robustness of our approach, we deliberately targeted proteins spanning a wide range of native expression levels. To correlate GFP fluorescence to protein abundance, we used a published ribosome profiling dataset from 293T cells as a reference for protein expression levels (25). Ribosome profiling is a high-throughput sequencing-based method that measures the density of ribosomes present on cellular mRNAs, thus providing a measure of protein synthesis rate (26). For each gene, ribosome density as measured by ribosome profiling is represented by a Reads Per Kilobase of transcript per Million mapped reads (RPKM) value. Because the abundance of a given protein is closely associated with the rate of its synthesis, RPKM data is a reasonable proxy for absolute protein expression levels (27). The relationship between flow cytometry GFP signal of knock-in cells and RPKM level for all 48 proteins we tested is shown in Figure 2B. GFP fluorescence intensity and predicted protein abundance for the 30 positive knock-in lines are well correlated (blue dots on Fig. 2B), indicating that GFP11 expression reports on the native expression level of the target protein. To estimate a minimal expression level compatible with GFP detection by flow cytometry, we found that an expression level of 27 RPKM would yield a GFP signal 2 standard deviations above background fluorescence (light blue line on Fig. 2B) based on a regression from our data (solid line on Fig. 2B). In the ribosome profiling dataset, about 30% of proteins expressed in 293T cells are found above this 27 RPKM threshold (defining here a protein as expressed if its RPKM is non-zero). In other words, this qualitative analysis suggests that ∼30% of proteins in a given cell line have an expression level compatible with the detection of GFP11 knock-in cells by flow cytometry.

Low protein expression is likely the main determinant for the lack of GFP-positive cells detected by flow cytometry in 37% of the genes we targeted. Comparing expression levels of the successful vs. unsuccessful sets of targets revealed that unsuccessful targets have significantly lower expression levels (median expression: 180 vs. 40 RPKM, respectively; see box plots in Fig. 2B). Therefore, the fluorescent signal for some of these failed targets might simply be below the detection limit of our flow cytometry assay. This is exemplified by NUP35 (Fig. 2C), a nuclear-pore complex protein of low expression level (43 RPKM). NUP35 GFP11-tagged cells scored negative by flow cytometry, but confocal microscopy analysis revealed cells exhibiting dim GFP fluorescence clearly restricted to foci on the nuclear membrane (Fig. 2C), indicative of specific NUP35 tagging. Fluorescent detection of NUP35 is facilitated by the fact that NUP35 concentrates in specific foci so that proteins of similar abundance but with a more diffuse localization pattern might be very hard to detect, even by microscopy. Altogether, our data shows that relying on endogenous expression levels poses a particular challenge for the study of low-abundance proteins, which in fact make up the bulk of proteins in human cells.

**A scalable strategy for the knock-in of GFP11 repeats enables fluorescent detection of low-expression proteins.** Our results highlight the difficulty in studying proteins of low abundance while maintaining native expression levels. How can these two elements be reconciled? As we have previously shown (19), the GFP11 system offers an elegant solution: by tagging a protein with repeats of the GFP11 sequence, multiple GFP1–10 fragments can be recruited to the same polypeptide thereby increasing the fluorescent signal of the target (Fig. 3B). Importantly, tagging with GFP11 repeats preserves native protein function. For example, the tandem arrangement of 7 GFP11 sequences enabled us to readily track a single transport particle in primary cilia without affecting its motility (19).

**Fig. 3.**
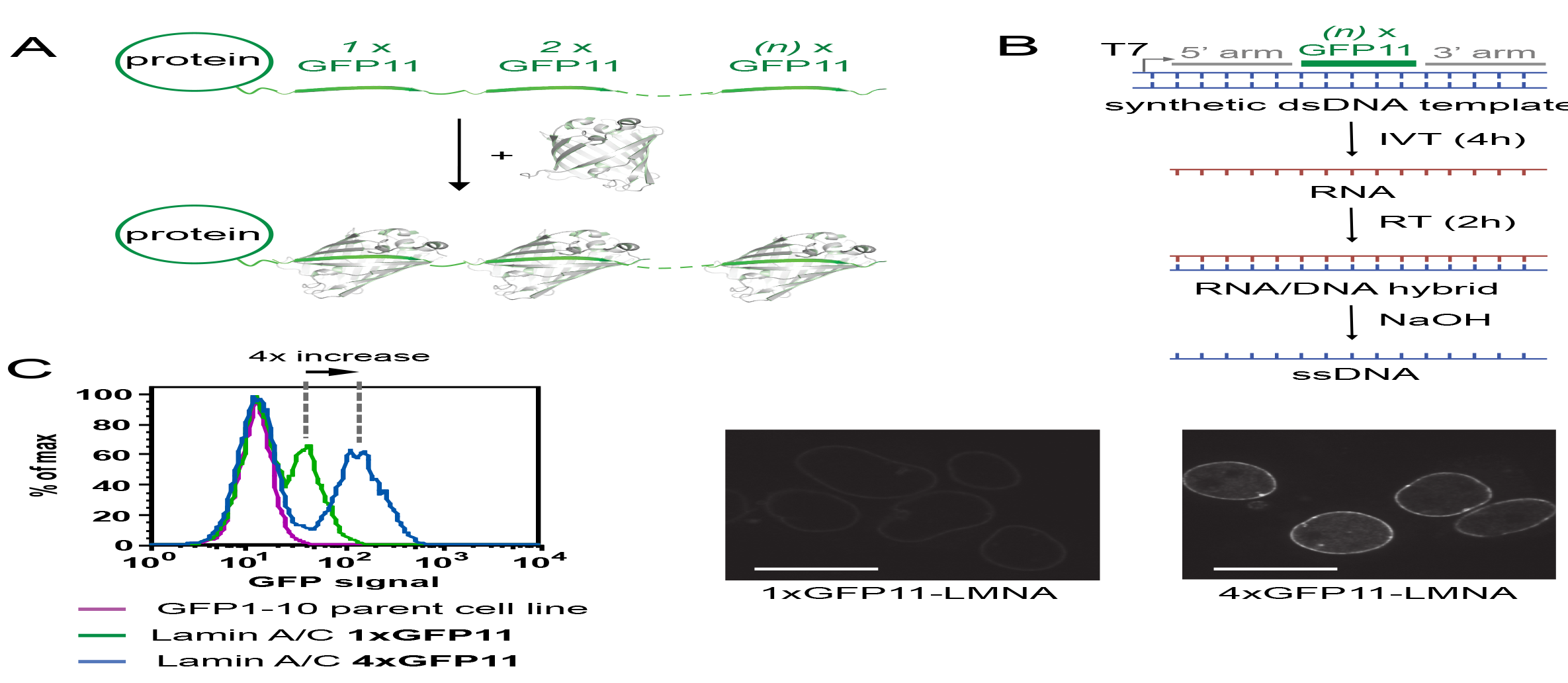
Knock-in of GFP11 repeats increases GFP fluorescence. (A) Principle of fluorescent tagging with GFP11 repeats. (B) Experimental worflow for ssDNA synthesis of HDR templates. See text for details. IVT: in vitro transcription. RT: reverse transcription. (C) Comparison of 1xGFP11 vs. 4xGFP11 knock-in at the Lamin A/ C N-terminus as analyzed by flow cytometry (left panel) and confocal microscopy (GFP fluorescence, right panels; scale bars =10 μm). Microscopy images were taken under identical exposure conditions and are shown using identical brightness and contrast settings, and can therefore be directly compared to one another.

We sought to develop an experimental strategy that would allow knock-in of GFP11 repeats while preserving the scalability, specificity and efficiency of our protocols. In particular, we reasoned that using a ssDNA form of HDR template would be advantageous since ssDNA donors have been shown to be more efficient and less prone to non-specific integration than their double-stranded counterparts (14, 28). Because GFP11 repeats exceed the current size limitation for ssDNA synthesis, we exploited the availability of large synthetic double-stranded DNA fragments for the production of ssDNA templates by adapting a method originally described for the synthesis of imaging probes (29). Our strategy starts with a synthetic (commercial) dsDNA fragment containing a T7 promoter followed by a cassette of GFP11–repeats flanked by homology arms (Fig. 3B). T7 in-vitro transcription followed by reverse transcription yields a DNA:RNA hybrid product. The RNA strand can be readily hydrolyzed at high pH to produce a corresponding ssDNA molecule (Fig. 3B). By using SPRI magnetic beads for all purification steps, these protocols can be carried out in multi-well format and in less than 8 hours. Together with the wide availability of commercial resources for synthetic dsDNA synthesis, our method enables the fast and scalable production of ssDNA HDR templates irrespective of sequence length.

To evaluate this approach, we prepared a ssDNA template for the tagging of the lamin A/C N-terminus with 4 repeats of GFP11 (including ∼300 nt homology arms flanking a 4xGFP11 tagging cassette of ∼250 nt). Flow cytometry analysis (Fig. 3C) revealed that the 4xGFP11 cassette was integrated with similar efficiency to the 1xGFP11 counterpart. In addition, 4xGFP11 tagging led to a corresponding 4-fold increase in fluorescence intensity (Fig. 3C). This increase is also apparent in microscopy images taken using identical exposure levels (Fig. 3C, right panels). This microscopy analysis also confirmed that GFP signal is limited to the inner nuclear membrane, confirming knock-in specificity. Altogether, these results validate our experimental strategy for the scalable and high-efficiency tagging of endogenous loci with GFP11 repeats. By lowering the expression level required for detection, GFP11 repeats enable the study of low-abundance proteins in their native cellular context. These methods pave the way for the construction of GFP-tagged cell libraries covering a majority of the human proteome. For example, while the analysis above indicated that only about 30% the proteome is accessible with a single GFP11 (RPKM>27), about 60% of all expressed proteins could be detected with 4xGFP11 repeats (assuming a four-fold lower detection limit, i.e. RPKM>6.8).

**Isolation of native protein complexes from GFP11 knock-in cells.** One of the great advantages of GFP is its versatility as both a fluorescent marker and a very effective handle for the immuno-purification of native complexes (16). The use of anti-GFP pull-downs for the high-resolution mapping of protein interactions by mass spectrometry is illustrated by recent studies using human lines containing GFP-tagged genes expressed on bacterial artificial chromosomes (17, 18). Therefore, we envisioned that GFP11 endogenous knock-in cells lines might be a valuable resource for the study of native protein-protein interactions in human cells.

We first confirmed that the non-covalent GFP11/GFP1-10 assembly can be efficiently captured by conventional anti-GFP reagents. We focused on four well-established multi-protein complexes: cohesin (30), the SEC61 translocon (31), clathrin (32) and the SPOTS sphingolipid synthesis complex (33). For each, we tagged a single subunit in 293T^GFP1–10^ cells, FACS-sorted knock-in cells and prepared lysates that were incubated with a commercial anti-GFP nanobody resin. After extensive washing of the resin, we eluted bound proteins by denaturation in SDS buffer and analyzed protein complexes by Western blot. For all four complexes we were able to recover the GFP11-tagged bait as well as its expected interaction partners (Fig. 4A). Because bound proteins can be directly digested on-beads and affinity capture is sufficient for quantitative mass-spectrometry experiments (17, 18), our results demonstrate the utility of endogenous GFP11 knock-in for the proteomic analysis of native protein complexes.

**Fig. 4.**
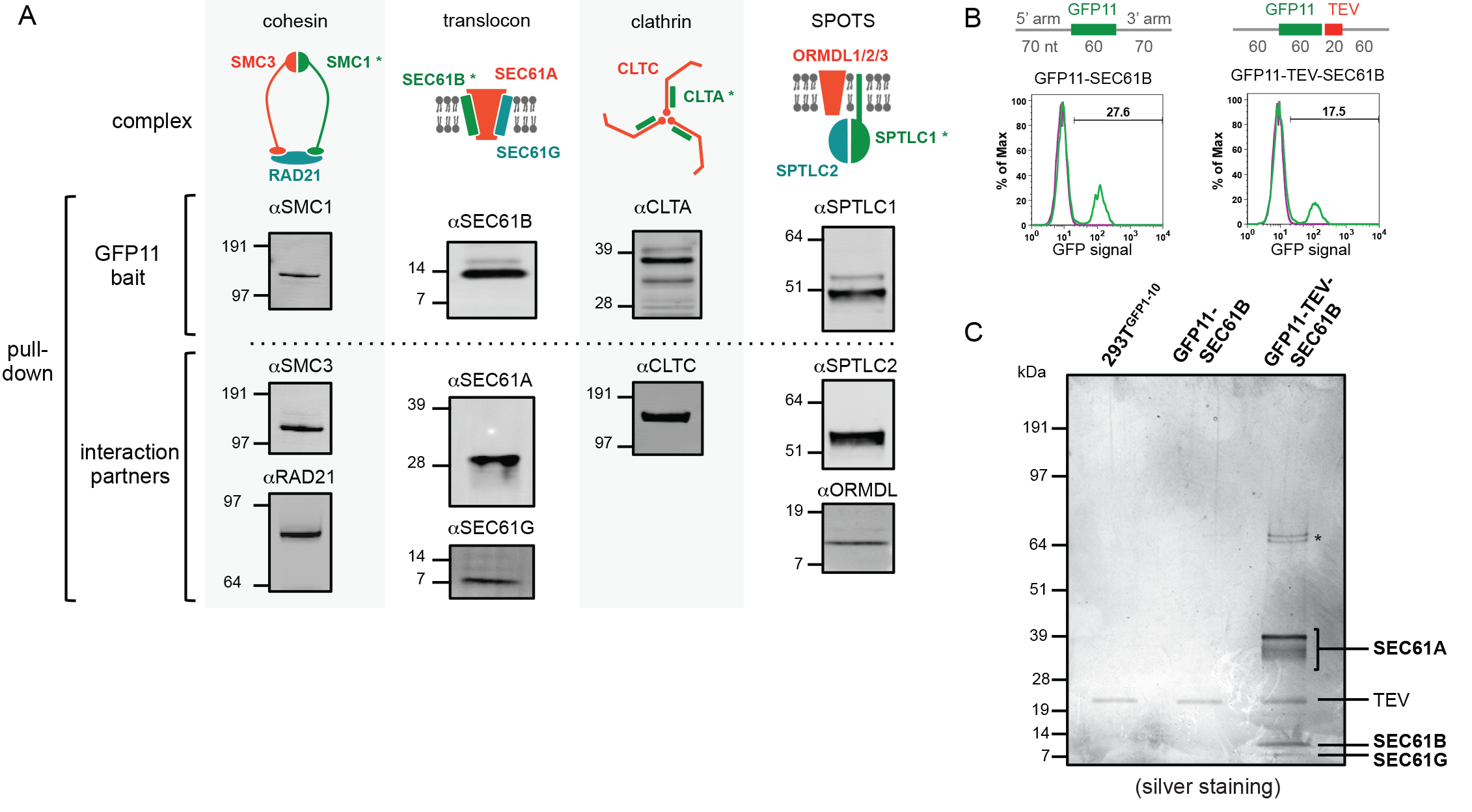
Isolation of native protein complexes in GFP11 knock-in cells by GFP immuno-precipitation. (A) Western-blot analysis following GFP immuno-precipitation. Four distinct protein complexes (cohesion, translocon, clathrin and SPOTS) were studied. For each complex, a single subunit was tagged with GFP11 (”GFP11 bait“,marked by an asterisk in corresponding drawings). Proteins were captured on anti-GFP resin, washed extensively and eluted in SDS buffer. Protein content was analyzed by SDS-PAGE and Western blot using protein-specific antibodies. Both GFP11 bait and expected interaction partners can be recovered. Numbers represent the migration of molecular weight markers (in kDa). (B) Comparison of knock-in efficiency of GFP11 vs. GFP11-TEV tag sequences at the SEC61B N-terminus, as analyzed by flow cytometry. Corresponding ssDNA HDR templates are shown. (C) Recovery of purified SEC61 complex following on-resin TEV cleavage. Proteins were captured on anti-GFP resin, washed extensively and eluted by incubation with TEV protease. Eluates were analyzed by SDS-PAGE and silver staining. SEC61 proteins are marked, as well as unidentified interaction partners (asterisk) and TEV protease.

For applications in which the recovery of purified proteins is advantageous (e.g., activity assays or structural studies), we modified our tagging cassette to include a TEV site to allow the specific release of captured proteins by protease treatment. To pilot this approach, we tagged the SEC61B N-terminus with GFP11 followed by a TEV recognition sequence. Because the TEV recognition sequence is short enough (7 amino acids), the GFP11-TEV cassette can be included on a 200–nt synthetic ssDNA oligo template (Fig. 4B). Knock-in efficiencies of GFP11-TEV vs. GFP11 alone were comparable (respectively 18% and 28%, Fig. 4B). We FACS-sorted knock-in cells, captured tagged proteins on anti-GFP beads and eluted by treatment with TEV protease. Analysis of the eluate by SDS-PAGE and silver staining (Fig. 4C) showed the specific elution of the entire SEC61 complex (SEC61A, SEC61B and SEC61G, see Fig. S4), together with unidentified interaction partners (marked by an asterisk in Fig. 4C). The comprehensive analysis of the SEC61 interactome is beyond the scope of the present study, but this pilot experiment demonstrates that our tagging method can be easily adapted to include protease recognition sites for the release of captured proteins. In particular, this purification strategy yields very pure material despite the low abundance of endogenous proteins: no background staining was detected in control samples using lysates from either the GFP1–10 parent cell line or a GFP11–SEC61B construct that does not include a TEV recognition sequence (Fig. 4C). These controls also demonstrate the high specificity of anti-GFP nanobody reagents for the capture of tagged proteins.

## DISCUSSION

Altogether, our results establish GFP11 RNP knock-in as a powerful strategy for the fast and efficient generation of endogenously tagged human cell lines. Our approach has several key advantages. First, contrary to designs that require the multi-step preparation of HDR targeting vectors, all the protocols we describe require no molecular cloning and can be carried out very rapidly and in large-scale format. Second, Cas9 RNP electroporation and ssDNA templates enable very high knock-in efficiency while minimizing off-target cleavage or non-specific tag integration (14). Third, the GFP11 system provides a simple solution for the study of low-abundance proteins because knock-in of GFP11 repeats increases fluorescence signal. Fourth, GFP is a particularly versatile tool that enables the study of both protein localization and protein-protein interactions. Finally, the utility of endogenously tagged cell lines is evident, allowing the function of a protein to be characterized under the control of native regulators of gene expression and without disturbing endogenous interaction stoichiometry. In this respect, the small size of the GFP11 cassette is advantageous because its introduction into a locus of interest is relatively seamless, minimizing perturbation of the surrounding genomic structure. Together, the methods presented here provide scalability, specificity, versatility and selectability and pave the way for the genome-scale construction of human cell lines tagged with GFP at endogenous loci. Interestingly, we recently described a split-sfCherry construct using a design similar to the GFP11 system (19). All our protocols can be directly adapted to any other split-fluorescent proteins, enabling the construction of multi-color tagged cell lines. Furthermore, other functional sequences can be coupled to GFP11 to tag proteins for various applications (e.g. protease sites for elution, or degron sequences for the specific control of protein expression (34)) and our results with 4xGFP11 knock-in show that long tagging cassettes can be integrated with high efficiency. Lastly, GFP11-tagged cell lines could be a valuable resource for structural genomics efforts. Indeed, GFP tagging is a powerful tool to identify biochemically stable protein complexes by fluorescent size-exclusion chromatography (35) and also enables the recovery of high-purity material suitable for structural characterization (especially by cryo-electron microscopy, which does not require large amounts of material).

Our approach also has a few limitations that should be addressed. The main restriction is the requirement for GFP1–10 expression in the cell line of interest. Here we used lentiviral methods for the integration of a GFP1–10 expression cassette for practicability. A more controlled strategy would be to insert the GFP1–10 cassette in an established safe harbor locus, where insertion of exogenous sequences is known to preserve genomic integrity (36). Safe harbor integration can be easily achieved, for example at the human AAVS1 locus (36). The cytoplasmic form of GFP1–10 can only complement with GFP11 accessible from the cytoplasm or the nucleus. To address this restriction, we have previously demonstrated that adding localization signals to GFP1-10 enables the labeling of GFP11–tagged proteins in other cellular compartments, such as using endoplasmic reticulum-localized GFP1–10 to label ER lumen proteins and extracellular domains of transmembrane proteins (19). A last limitation of our approach is inherent to any effort of protein tagging. It is possible that, in a subset of proteins, introduction of GFP11 would disturb protein function (for example by changing protein structure or shielding an important interaction interface). We believe that the small size of GFP11 is beneficial in this respect, as it should not affect much the native folding of the target protein. Importantly, GFP11 can be introduced interchangeably at either N-or C-terminus (or in any loop region) of a protein target, and it is likely that in cases where introducing the tag at one site is problematic, introducing it at another position would be well tolerated.

Finally, our strategy is also limited by any shortcomings of the CRISPR/Cas9 system. In particular, knock-in efficiency depends critically on the activity of the sgRNA used for genomic cleavage. Different sgRNA sequences can vary widely in term of potency, and prediction algorithms have been developed to overcome this issue (37, 38). But since HDR knock-in requires genomic cleavage close to the site of tag integration, for some genes the choice of sgRNAs to pick from might be scarce. However, our results are very encouraging in this respect. In our 48–gene library-scale experiment, we only tested a single sgRNA for each gene and saw a high rate (63%) of successful tagging. Alternatively, tagging a given protein at another site in the protein sequence might allow more optimal genomic cleavage. A last limitation is that, because 100% knock-in efficiency is not currently attainable, most targeted cells have only a single allele tagged. Moreover, because non-homologous-end joining is usually more prevalent than homologous-directed repair following Cas9 cleavage (14), it is likely that in some cells the non-tagged allele (or alleles, in polyploid cells) will contain indel mutations. We believe that, in most cases, this should not compromise the proper functional characterization of the target protein. In particular, working with polyclonal populations and using population averages helps mitigate the possible defects present in a small number of individual cells. Alternatively, single clones can be isolated to identify homozygous knock-in cells. The very high knock-in efficiencies that we report will significantly facilitate the successful isolation of homozygous clones.

Altogether, the results of our library-scale experiment highlight the applicability of GFP11 knock-in for the tagging of a large fraction of the human proteome. We anticipate that low expression level of a target protein will be an obstacle to the detection and selection of a subset of GFP11-tagged cells. The tagging of genes with GFP11 repeats provides a direct solution to this drawback. Notably, tagging with GFP11 repeats is not substantially more challenging than tagging with a single GFP11 sequence. Our protocols for the production of long ssDNA templates are simple, fast (<1 day) and cloning-free. Furthermore, the example of Lamin A/C tagging (Fig. 3C) demonstrates that 1xGFP11 and 4xGFP11 cassettes are integrated with comparable efficiency. Therefore, tagging with GFP11 repeats should be preferred for proteins expected to be expressed at low levels. On the other hand, for a small subset of targets we could not detect GFP positive cells despite their high predicted expression (Fig. 2B), suggesting that expression level is not the sole determinant for successful tagging. In some cases, this lack of detectable tagging might indicate that the Cas9/sgRNA complex failed to access and cut the target genomic sequence (for example, we have recently shown that high nucleosome occupancy can impede Cas9 access to DNA (39)). As a solution, tagging could be achieved by using sgRNAs targeting alternative sites within the desired locus. In some other cases, the lack of GFP detection could originate from the lack of physical accessibility to the GFP11 tag for complementation with GFP1–10 (for example, if GFP11 is buried inside a structural pocket within the target protein). Then, introducing a longer linker between the target protein and the GFP11 tag would be beneficial.

Overall, we believe that the many advantages of GFP11 RNP knock-in far outweigh its potential limitations, especially for studies requiring the tagging of many different genes in parallel given the speed and scalability of our protocols. In addition, our protocols will directly benefit from the continued and rapid optimization of CRISPR/Cas9-based methods. Altogether, the experimental approach described here directly paves the way for the generation of genome-wide libraries of human cells harboring GFP-tagged proteins at their endogenous loci. This opens tremendous opportunities for the comprehensive characterization of the human proteome in a native cellular context.

## MATERIALS AND METHODS

**Nucleic acid reagents.** All synthetic nucleic acid reagents were purchased from Integrative DNA Technologies (IDT DNA, Coralville, IA). For knock-in of a single GFP11 sequence, 200–mer HDR templates were ordered in ssDNA form (Ultramer oligos). For knock-in of GFP11 repeats, HDR template was ordered in dsDNA form (gBlock fragments) and processed to ssDNA as described below. The complete set of DNA sequences used for the experiments described here can be found in Supplementary Dataset 1.

**293T^GFP1–10^generation and cell culture.** HEK 293T cells were cultured in high-glucose DMEM supplemented with 10% FBS, 1 mM glutamine and 100 ng/mL penicillin/streptomycin (Gibco). 293T^GFP1–10^ cells were generated by lentiviral integration from the vector pHR-SFFV-GFP1–10 described in (19) and a clonal cell line was isolated and used for knock-in experiments. Cells were maintained below 80% confluency.

**sgRNA in vitro transcription.** sgRNAs were prepared following methods by Lin et al. (14) with some modifications. sgRNAs were obtained by in vitro transcription of a DNA template of the following sequence: 5'– TAA TAC GAC TCA CTA TAG GNN NNN NNN NNN NNN NNN NNG TTT AAG AGC TAT GCT GGA AAC AGC ATA GCA AGT TTA AAT AAG GCT AGT CCG TTA TCA ACT TGA AAA AGT GGC ACC GAG TCG GTG CTT TTT TT-3' containing a T7 promoter (TAATACGACTCACTATAG), a gene-specific ∼20–nt sgRNA sequence starting with a G for optimal T7 transcription (GNNNNNNNNNNNNNNNNNNN) and a common sgRNA constant region. The DNA template was generated by overlapping PCR using a set of 4 primers: 3 primers common to all reactions (forward primer T25: 5'– TAA TAC GAC TCA CTA TAG–3'; reverse primer BS7: 5'– AAA AAA AGC ACC GAC TCG GTG C–3' and reverse primer ML611: 5'– AAA AAA AGC ACC GAC TCG GTG CCA CTT TTT CAA GTT GAT AAC GGA CTA GCC TTA TTT AAA CTT GCT ATG CTG TTT CCA GCA TAG CTC TTA AAC-3') and one gene-specific primer (forward primer 5'– TAA TAC GAC TCA CTA TAG GNN NNN NNN NNN NNN NNN NNG TTT AAG AGC TAT GCT GGA A–3'). For each template a 100μL PCR was set using iProof High-Fidelity Master Mix (Bio-Rad) reagents supplemented with 1 μM T25, 1 μM BS7, 20 nM ML611 and 20 nM gene-specific primer. The thermocycler setting consisted of: 95°C for 30 s, 30 cycles of {95°C for 15 s, 57°C for 15 s, 72°C for 15 s}, 72 °C for 30 s. The PCR product was purified on DNA Clean and Concentrator-5 columns (Zymo Research) following the manufacturer's instructions and eluted in 12 μL of RNAse-free DNA buffer (2 mM Tris pH 8.0 in DEPC-treated H2O). Next, a 100μL in vitro transcription reaction was set using 300 ng DNA template and 1000 U of T7 RNA polymerase in buffer containing (in mM):

40 Tris pH 7.9, 20 MgCl2, 5 DTT, 2 spermidine and 2 each NTP (New England BioLabs). Following a 4–h incubation at 37°C, the sgRNA product was purified on RNA Clean and Concentrator–5 columns (Zymo 14 Research) and eluted in 15 μL of RNAse-free RNA buffer (10 mM Tris pH 7.0 in DEPC-treated H2O). sgRNA quality was routinely checked by running 3 pg of the purified sgRNA on a 10% polyacrylamide gel containing 7M urea (Novex TBE-Urea gels, ThermoFisher Scientific).

**RNP assembly and electroporation.** Cas9/sgRNA RNP complexes were prepared following methods by Lin et al. (14) with some modifications. Cas9 protein (pMJ915 construct, containing two nuclear localization sequences) was expressed in E. coli and purified by the UC Berkeley Macrolab following protocols described by Jinek et al. (22). 293T^GFP1–10^ cells were treated with 200 ng/mL nocodazole (Sigma) for 15 hours before electroporation to increase HDR efficiency as shown by Lin et al. (14). RNP complexes were assembled with 100 pmol Cas9 protein and 130 pmol sgRNA just prior to electroporation, and combined with HDR template in a final volume of 10 μL. First, 130 pmol purified sgRNA was diluted to 6.5 μL in Cas9 buffer (final concentrations: 150 mM KCl, 20 mM Tris pH 7.5, 1 mM TCEP-HCl, 1 mM MgCl2, 10% v/v glycerol) and incubated at 70°C for 5 min. 2.5 μL of Cas9 protein (40 μM stock in Cas9 buffer, ie. 100 pmol) was then added and RNP assembly was carried out at 37°C for 10 min. Finally, 1 μL of HDR template (100 μM stock in Cas9 buffer, ie. 100 pmol) was added to this RNP solution. Electroporation was carried out in Amaxa 96–well shuttle Nuleofector device (Lonza) using SF-cell line reagents (Lonza) following the manufacturer's instructions. Nocodazole-treated 293T^GFP1–10^ cells were washed with PBS and resuspended to 104 cellsμL in SF solution immediately prior to electroporation. For each sample, 20 μL of cells (ie. 2×10^5^ cells) was added to the 10 μL RNP/template mixture. Cells were immediately electroporated using the CM130 program and transferred to 1 mL supplemented DMEM in a 24–well plate. Electroporated cells were cultured for 5 days prior to analysis.

**Preparation of 4×GFP11-LMNA ssDNA template**. 4×GFP11-LMNA ssDNA template was prepared from a commercial dsDNA fragment (gBlock, IDT DNA) containing the template sequence preceded by a T7 promoter, adapting a strategy first described by Chen et al. (29). The dsDNA fragment was first amplified by PCR (forward primer ML888: 5'– AGC TGA TAA TAC GAC TCA CTA TAG GG–3', reverse primer ML904: 5'– CGA CTT TCG CGC CAC TCA AGC −3') using Kapa HiFi reagents (Kapa Biosystems) in a 100–μL reaction containing 0.25 μM each primer, 10 ng DNA template and 0.3 mM dNTPs. Amplified dsDNA was purified using SPRI beads (AMPure XP resin, Beckman Coulter) at a 1:1 DNA:resin volume ratio (following manufacturer's instructions) and eluted in 25 μL RNAse-free H2O. Next, RNA was formed by T7 in vitro transcription using T7 HiScribe reagents (New England BioLabs) in a 50–μL reaction containing: 5 pmol dsDNA template, 10 mM each NTP and 5 μL HiScribe T7 polymerase. Following a 4–h incubation at 37°C, the reaction was treated with 4U TURBO DNAse (ThermoFisher Scientific) and incubated another 15 min at 37°C. The RNA product was then purified using SPRI beads at a 1:1 RNA:resin volume ratio and eluted in 60 μL RNAse-free H2O. DNA:RNA hybrid was then synthesized by reverse transcription using Maxima H RT reagents (ThermoFisher Scientific). First, a 42–μL solution (in nuclease-free water) containing 500 pmol RNA template, 1 nmol ML904 primer and 2.4 mM each dNTPs was incubated 5 min at 65°C and transferred on ice for 5 min to allow for primer annealing. 12 μL 5x Maxima buffer, 3 μL Maxima RT enzyme and 3 μL SUPERase In RNAse inhibitor were then added and the RT reaction was carried out for 45 min at 50°C. Finally, the RNA strand was hydrolyzed by the addition of 24 μL of NaOH solution (0.5 M NaOH + 0.25 M EDTA, in H2O) followed by incubation at 95°C for 10 min. The final ssDNA product was purified using SPRI beads at a 1:1.2 DNA:resin volume ratio and eluted in 15 μL H^2^O.

**Flow cytometry and analysis.** Analytical flow cytometry was carried out on a LSR II instrument (BD Biosciences) and cell sorting on a FACSAria II (BD Biosciences). Flow cytometry data analysis and figure preparation was done using the FlowJo software (FlowJo LLC). For the measurement of GFP signals in Figure 3B, flow cytometry traces were fitted with two Gaussian functions (the first Gaussian corresponding to background fluorescence, the second Gaussian to specific GFP fluorescence). GFP signal is measured by the difference: (average specific GFP fluorescence) -(average background fluorescence). Double Gaussian fit was particularly important to measure GFP signal of low-expression proteins, for which background and specific GFP signals have significant overlap (ex: SPTLC1 in Figure 2B).

**Protein pull-down.** For each sample, the cell pellet from a 15–cm plate culture was resuspended in 1.5 mL GFP buffer (150 mM K-acetate, 50 mM Hepes pH 6.8, 2 mM MgCl2, 1 mM CaCl2, 15% v/v glycerol) supplemented with 1.5% w/v digitonin (high purity, Merck Millipore) and protease inhibitors (cOmplete EDTA-free cocktail, Roche), and incubated 2 h at 4°C, rotating. The lysate was then clarified by centrifugation (20,000xg, 30 min, 4°C) and the supernatant incubated with 8 μL anti-GFP resin slurry (GFP-Trap_A resin, ChromoTek) for 2 h at 4°C, rotating. The resin was then washed 3 times with wash buffer (GFP buffer + 0.1 % digitonin). For Western blot analysis, proteins were eluted by boiling the washed resin in SDS buffer (50 mM Tris pH 6.8, 2% w/v SDS, 1% β-ME, 6% glycerol; final concentrations). For TEV elution, the washed resin was incubated with 0.5 μg of His6-TEV protease (Sigma) overnight at 4°C.

**Primary antibodies used for Western-blot.** Anti-SMC1: ProMab 20426. Anti-SMC3: Abcam ab9263. Anti-RAD21: Abcam ab992. Anti-SEC61B: Cell Signaling Technologies D5Q1W. Anti-SEC61A: Cell Signaling Technologies D7Q6V. Anti-SEC61G: Proteintech 11147–2–AP. Anti-CLTA: X16, gift from Yvette Schollmeier, F. Brodsky lab. Anti-CLTC: Santa Cruz Biotechnology sc–12734. Anti-SPTLC1: BD Biosciences 611305. Anti-SPTLC2: ProSci 6305. Anti-ORMDL: Abcam ab128660. All antibodies were used at 1:1000 dilution.

**Imaging.** Cells were grown in 96-well glass bottom plates with #1.5 high performance cover glass (In Vitro Scientific) coated with Fibronectin (Roche) for 48 hours then fixed with 4% Paraformaldehyde (Electron Microscopy Sciences, Cat. #15710-S) for 15 minutes at room temperature. The fixed cells were imaged on an inverted Nikon Ti-E microscope, Yokogawa CSU-22 confocal scanner unit, Plan Fluor 10x/0.3 NA objective or Plan Apo VC 60x/1.4 NA oil objective, an Andor EM-CCD camera (iXon DU897) and Micro-Manager software. All imaging experiments were performed at UCSF Nikon Image Center. For the comparison of 1xGFP11-LMNA and 4xGFP11-LMNA in Figure 3C, exactly same excitation power, exposure time and brightness and contrast were used. The brightness and contrast for other images were automatically set by ImageJ. For immunocytochemistry, mouse monoclonal anti-histone H2B ([1:50] abcam, ab52484) antibody, rabbit polyclonal antibodies anti-lamin A/C ([1:20] Santa Cruz Biotechnology, H110), anti-cAMP protein kinase catalytic subunit ([1:1000] abcam, ab26322), and anti-CBX/HP1 beta ([1:100] abcam, ab10478) were used. Anti-mouse or anti-rabbit Donkey secondary antibodies (Jackson Immuno Research Laboratories, INC.) were conjugated with Alexa Fluor 647 or Cy5, respectively. The fixed cells were permeabilized with 0.1% Triton X-100 (Sigma), blocked with 5% BSA (Jackson Immuno Research Laboratories, INC.) in PBS, stained with primary antibodies and secondary antibodies at 4 °C overnight.

## ACKNOWLEDGEMENTS

We thank B. Staahl and S. Lin in the J. Doudna laboratory (UC Berkeley) for advice with RNP preparation, E. Crawford in the J. DeRisi's laboratory (UCSF) for advice with sgRNA purification and A. Banfal in the Weissman laboratory for help with 4xGFP11-LMNA template preparation. M.D.L. is a fellow of the Jane Coffin Childs Memorial Funds for Medical Research. S.S. is supported by a Japan Society for the Promotion of Science Postdoctoral Fellowship for Overseas Researchers. This work was supported by NIH R21MH101688 (B.H. and D.K.), NIH Director's New Innovator Award DP20D008479 (B.H.), and the Howard Hughes Medical Institute (J.S.W.).

## FIGURE LEGENDS

**Fig. 1.Endogenous GFP11 tagging using Cas9 RNP.** (A) Principle of GFP11-mediated tagging. (B) Experimental workflow. T7 IVT: in vitro transcription using T7 polymerase. (C) GFP11 knock-in at the Lamin A/C N-terminus. Knock-in efficiency was analyzed by flow cytometry (left panel, showing the distribution of GFP fluorescence as histogram plot). GFP-positive cells were isolated by FACS and characterized by confocal microscopy (right panel, showing GFP fluorescence, scale bar = 10 μm).

**Fig. 2.Library-scale GFP11 tagging of 48 different gene targets.** (A) Examples of successful targets showing knock-in efficiency (flow cytometry histograms, top panels) and confocal microscopy analysis (GFP fluorescence, bottom panels; scale bar =10 μm). As GFP intensity varies widely across different targets, the different images showed here use different levels of brightness and contrast. (B) Correlation between target expression level (defined as ribosome profiling RPKM) and GFP signal (as measured by flow cytometry, arbitrary units scaled to background fluorescence =1). The 30 successful targets and 18 unsuccessful targets are shown as blue and brown dots, respectively. For successful targets, a linear regression is shown (solid line, Pearson R =0.69). Insert: box plots showing RPKM distribution for unsuccessful vs. successful targets. Boxes represent 25th, 50th and 75th percentiles. Whiskers represent minimum and maximum values. (C) Analysis of NUP35 GFP11 knock-in by flow cytometry (top panel) and confocal microscopy (bottom panel; scale bar =10 μm). NUP35 knock-in cells are not detected by flow cytometry but can be identified by microscopy.

**Fig. 3.Knock-in of GFP11 repeats increases GFP fluorescence.** (A) Principle of fluorescent tagging with GFP11 repeats. (B) Experimental workflow for ssDNA synthesis of HDR templates. See text for details. IVT: in vitro transcription. RT: reverse transcription. (C) Comparison of 1xGFP11 vs. 4xGFP11 knock-in at the Lamin A/C N-terminus as analyzed by flow cytometry (left panel) and confocal microscopy (GFP fluorescence, right panels; scale bars =10 μm). Microscopy images were taken under identical exposure conditions and are shown using identical brightness and contrast settings, and can therefore be directly compared to one another.

**Fig. 4.Isolation of native protein complexes in GFP11 knock-in cells by GFP immuno-precipitation.** (A)Western-blot analysis following GFP immuno-precipitation. Four distinct protein complexes (cohesin, translocon, clathrin and SPOTS) were studied. For each complex, a single subunit was tagged with GFP11 (“GFP11 bait“, marked by an asterisk in corresponding drawings). Proteins were captured on anti-GFP resin, washed extensively and eluted in SDS buffer. Protein content was analyzed by SDS-PAGE and Western blot using protein-specific antibodies. Both GFP11 bait and expected interaction partners can be recovered. Numbers represent the migration of molecular weight markers (in kDa). (B) Comparison of knock-in efficiency of GFP11 vs. GFP11-TEV tag sequences at the SEC61B N-terminus, as analyzed by flow cytometry. Corresponding ssDNA HDR templates are shown. (C) Recovery of purified SEC61 complex following on-resin TEV cleavage. Proteins were captured on anti-GFP resin, washed extensively and eluted by incubation with TEV protease. Eluates were analyzed by SDS-PAGE and silver staining. SEC61 proteins are marked, as well as unidentified interaction partners (asterisk).

**Suppl. Fig. S1.**
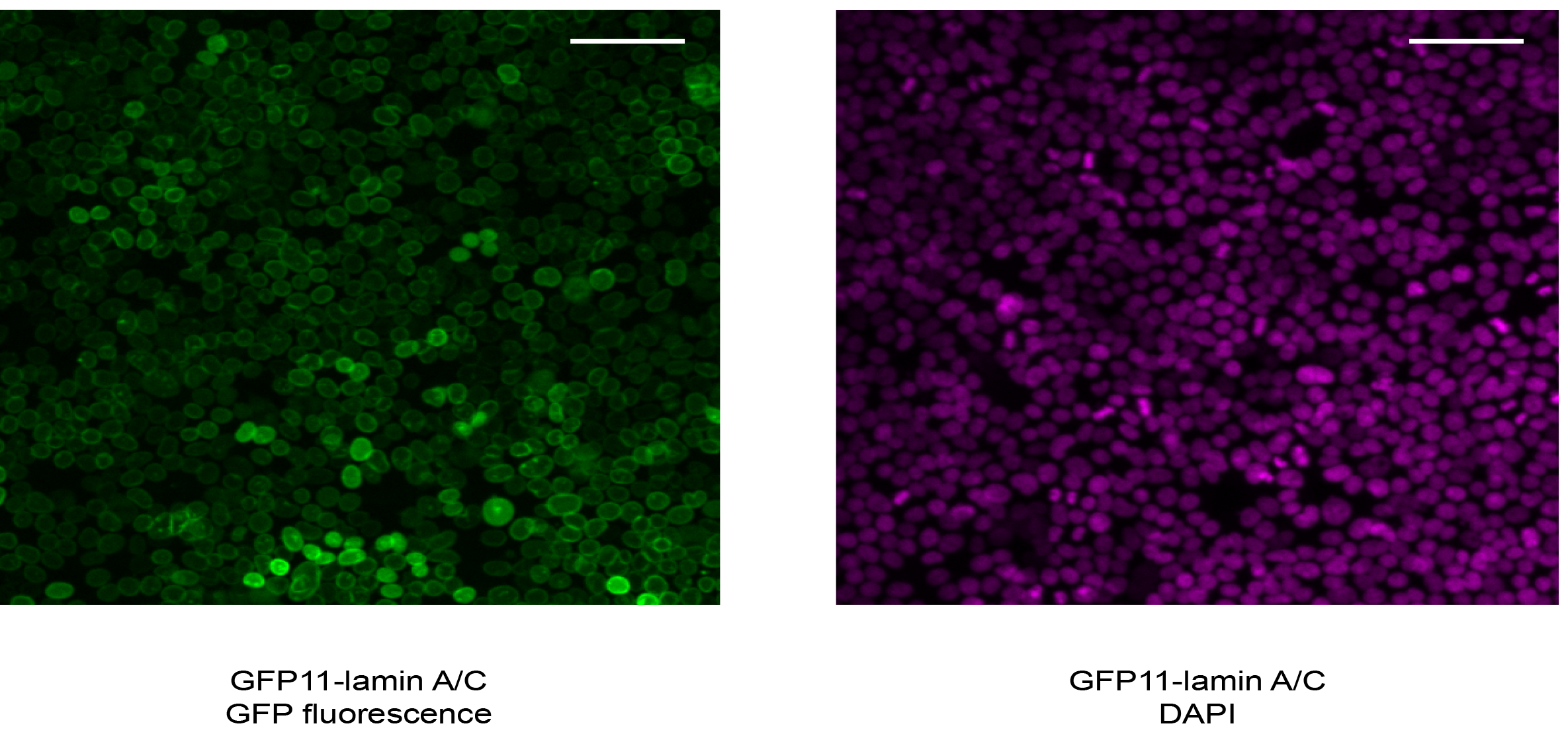
**Low-magnification microscopy analysis of GFP11-lamin A/C cells.**FACS-sorted GFP11-lamin A/C knock-in cells were stained with DAPI and analyzed by confocal microscopy. Scale bars =50 μm.

**Suppl. Fig. S2.**
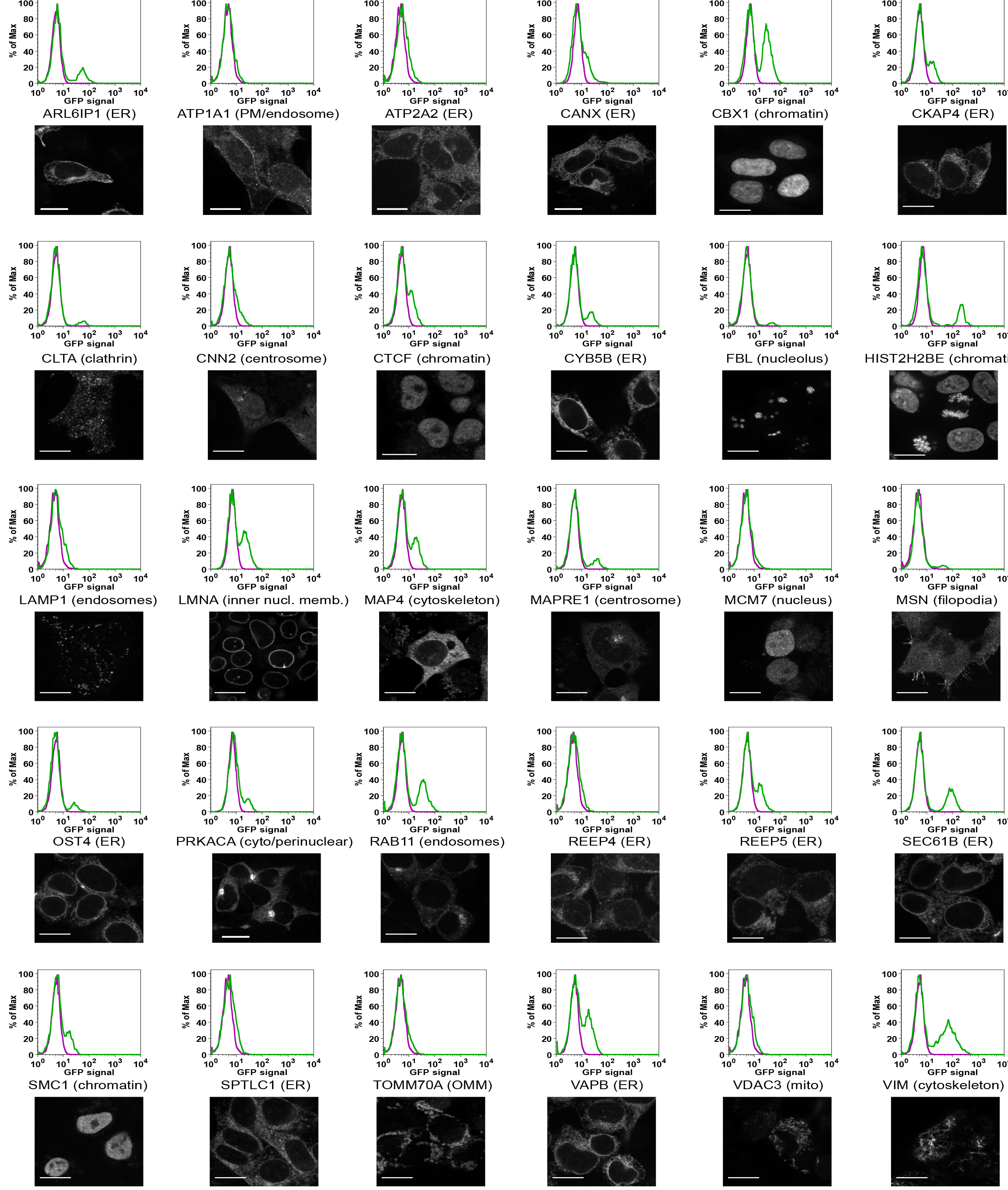
**Analysis of GFP-positive targets by flow cytometry and fluorescence microscopy.**All 30 gene targets succesfully tagged by GFP11 are shown. For each target, both flow cytometry analysis (top panels) and representative confocal microscopy pictures (bottom panels) are shown. Expected sub-cellular localization of each target is indicated (ER: endoplasmic reticulum, PM: plasma membrane, cyto: cytosol, OMM: outer mitochondrial membrane, mito: mitochondria). Scale bars =10 μm.

**Suppl. Fig. S3.**
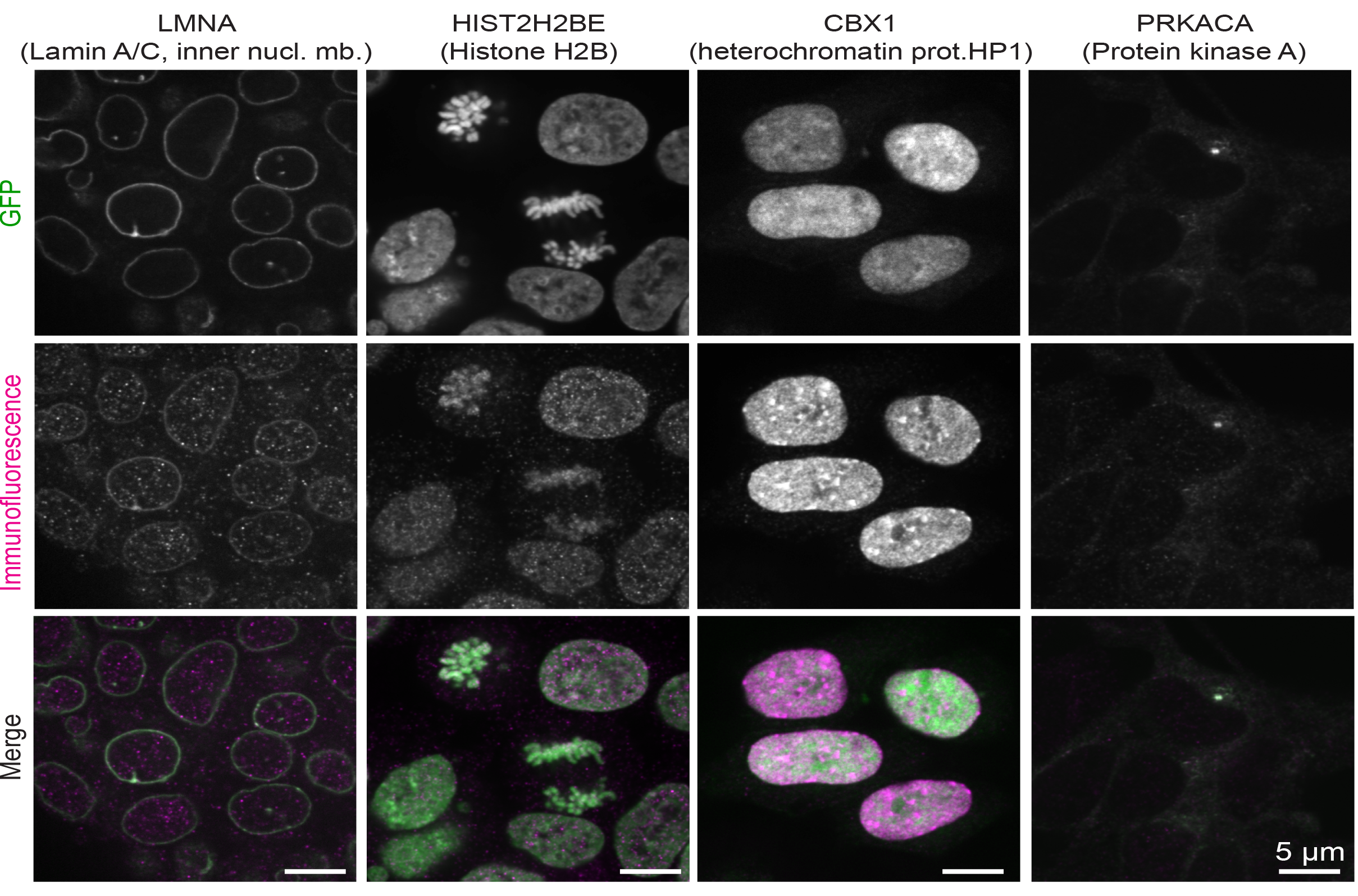
**Comparison of GFP signal and protein localization by immunofluorescence.**For the four knock-in targets shown, GFP-positive cells were FACS-sorted, fixed and stained with antibodies to the corresponding protein target. In each case, GFP fluorescence (top panels) and immuno-staining (middle panels) are compared. Scale bars: 5 μm.

**Suppl. Fig. S4.**
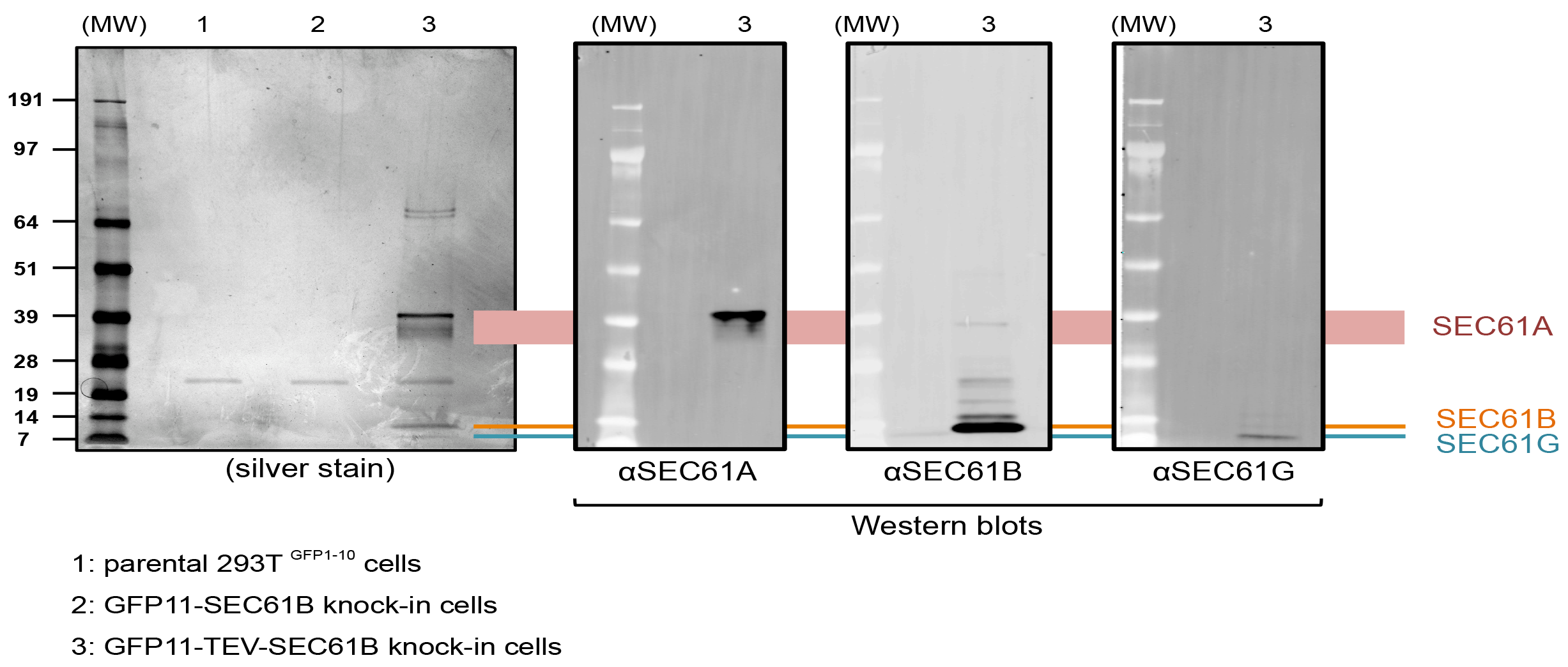
**Identification of SEC61 subunits by Western blot.** Related to Figure 4. To confirm the identity of proteins observed by silver staining in the GFP11-TEV-SEC61B eluate (lane 3), the eluate was analyzed by Western Blot using subunit-specific antibodies. All images were aligned according to the position of molecular-weight standards (MW). Subunit identity was assigned based on matching position between silver stain and Western Blot images.

## REFERENCES

1. International Human Genome Sequencing Consortium (2004)Finishing the euchromatic sequence of the human genome. Nature 431(7011):931–945.

2. Hanson AD, Pribat A, Waller JC, de Crécy-Lagard V (2010)'Unknown' proteins and “orphan” enzymes: the missing half of the engineering parts list”and how to find it. Biochem J 425(1):1–11.

3. Dey G, Jaimovich A, Collins SR, Seki A, Meyer T (2015)Systematic Discovery of Human Gene Function and Principles of Modular Organization through Phylogenetic Profiling. Cell Rep. doi:10.1016/j.celrep.2015.01.025.

4. Schuldiner M, Weissman JS (2013)The Contribution of Systematic Approaches to Characterizing the Proteins and Functions of the Endoplasmic Reticulum. Cold Spring Harb Perspect Biol. doi:10.1101/cshperspect.a013284.

5. Huh W-K, et al. (2003)Global analysis of protein localization in budding yeast. Nature 425(6959):686–691.

6. Gavin A-C, et al. (2002)Functional organization of the yeast proteome by systematic analysis of protein complexes. Nature 415(6868):141–147.

7. Krogan NJ, et al. (2006)Global landscape of protein complexes in the yeast Saccharomyces cerevisiae. Nature 440(7084):637–643.

8. Baudin A, Ozier-Kalogeropoulos O, Denouel A, Lacroute F, Cullin C (1993)A simple and efficient method for direct gene deletion in Saccharomyces cerevisiae. Nucleic Acids Res 21(14):3329–3330.

9. Ghaemmaghami S, et al. (2003)Global analysis of protein expression in yeast. Nature 425(6959):737–741.

10. Chong YT, et al. (2015)Yeast Proteome Dynamics from Single Cell Imaging and Automated Analysis. Cell 161(6):1413–1424.

11. Breker M, Schuldiner M (2014)The emergence of proteome-wide technologies: systematic analysis of proteins comes of age. Nat Rev Mol Cell Biol 15(7):453–464.

12. Cong L, et al. (2013)Multiplex genome engineering using CRISPR/Cas systems. Science 339(6121):819–823.

13. Jinek M, et al. (2013)RNA-programmed genome editing in human cells. Elife 2:e00471.

14. Lin S, Staahl B, Alla RK, Doudna JA (2014)Enhanced homology-directed human genome engineering by controlled timing of CRISPR/Cas9 delivery. Elife 3. doi:10.7554/eLife.04766.

15. Kim S, Kim D, Cho SW, Kim J, Kim JS (2014)Highly efficient RNA-guided genome editing in human cells via delivery of purified Cas9 ribonucleoproteins. Genome Research 24(6):1012–1019.

16. Cristea IM, Williams R, Chait BT, Rout MP (2005)Fluorescent proteins as proteomic probes. Mol Cell Proteomics 4(12):1933–1941.

17. Hein MY, et al. (2015)A Human Interactome in Three Quantitative Dimensions Organized by Stoichiometries and Abundances. Cell 163(3):712–723.

18. Hubner NC, et al. (2010)Quantitative proteomics combined with BAC TransgeneOmics reveals in vivo protein interactions. J Cell Biol 189(4):739–754.

19. Kamiyama D, et al. (2016)Versatile protein tagging in cells with split fluorescent protein. Nat Commun 7:11046.

20. Cabantous S, Terwilliger TC, Waldo GS (2005)Protein tagging and detection with engineered self-assembling fragments of green fluorescent protein. Nat Biotechnol 23(1):102–107.

21. Kent KP, Childs W, Boxer SG (2008)Deconstructing green fluorescent protein. J Am Chem Soc 130(30):9664–9665.

22. Jinek M, et al. (2012)A programmable dual-RNA-guided DNA endonuclease in adaptive bacterial immunity. Science 337(6096):816–821.

23. Yang H, et al. (2013)One-Step Generation of Mice Carrying Reporter and Conditional Alleles by CRISPR/Cas-Mediated Genome Engineering. Cell 154(6):1370–1379.

24. DeAngelis MM, Wang DG, Hawkins TL (1995)Solid-phase reversible immobilization for the isolation of PCR products. Nucleic Acids Res 23(22):4742–4743.

25. Jan CH, Williams CC, Weissman JS (2014)Principles of ER cotranslational translocation revealed by proximity-specific ribosome profiling. Science 346(6210):1257521.

26. Ingolia NT, et al. (2012)The ribosome profiling strategy for monitoring translation in vivo by deep sequencing of ribosome-protected mRNA fragments. Nat Protoc 7(8):1534–1550.

27. Li G-W, Burkhardt D, Gross C, Weissman JS (2014)Quantifying absolute protein synthesis rates reveals principles underlying allocation of cellular resources. Cell 157(3):624–635.

28. Chen F, et al. (2011)High-frequency genome editing using ssDNA oligonucleotides with zinc-finger nucleases. Nat Methods 8(9):753–755.

29. Chen KH, Boettiger AN, Moffitt JR, Wang S, Zhuang X (2015)RNA imaging. Spatially resolved, highly multiplexed RNA profiling in single cells. Science 348(6233):aaa6090.

30. Peters J-M, Tedeschi A, Schmitz J (2008)The cohesin complex and its roles in chromosome biology. Genes Dev 22(22):3089–3114.

31. Park E, Rapoport TA (2012)Mechanisms of Sec61/SecY-mediated protein translocation across membranes. Annu Rev Biophys 41:21–40.

32. Brodsky FM (2012)Diversity of clathrin function: new tricks for an old protein. Annu Rev Cell Dev Biol 28:309–336.

33. Breslow DK, Weissman JS (2010)Membranes in balance: mechanisms of sphingolipid homeostasis. Mol Cell 40(2):267–279.

34. Nishimura K, Fukagawa T, Takisawa H, Kakimoto T, Kanemaki M (2009)An auxin-based degron system for the rapid depletion of proteins in nonplant cells. Nat Methods 6(12):917–922.

35. Kawate T, Gouaux E (2006)Fluorescence-detection size-exclusion chromatography for precrystallization screening of integral membrane proteins. Structure 14(4):673–681.

36. Sadelain M, Papapetrou EP, Bushman FD (2012)Safe harbours for the integration of new DNA in the human genome. Nat Rev Cancer 12(1):51–58.

37. Doench JG, et al. (2014)Rational design of highly active sgRNAs for CRISPR-Cas9-mediated gene inactivation. Nat Biotechnol 32(12):1262–1267.

38. Moreno-Mateos MA, et al. (2015)CRISPRscan: designing highly efficient sgRNAs for CRISPR-Cas9 targeting in vivo. Nat Methods 12(10):982–988.

39. Horlbeck MA, et al. (2016)Nucleosomes impede Cas9 access to DNA in vivo and in vitro. Elife 5. doi:10.7554/eLife.12677.

